# shRNA Glut-1 inhibits cell viability, apoptosis and migration of laryngeal carcinoma HEp-2 cells through regulating Beclin-1-mediated autophagy

**DOI:** 10.1101/2020.02.24.962449

**Authors:** Wen-Dong Wang, Jin-Long Zhu, Shui-Hong Zhou, Jun Fan, Yang-Yang Bao

## Abstract

**Objective:** Glut-1 is a key regulator in the process of glucose uptake. Previous studies have shown that Glut-1 affects autophagy. However, it is unclear whether there is a correlation between Glut-1 and autophagy in the progression of laryngeal carcinoma. This study was performed to investigate the role of Glut-1 in the development of laryngeal carcinoma.

**Methods:** A stable HEp-2 cell model was constructed by Glut-1 and Beclin-1 shRNA lentiviral infection. The autophagosome was measured by transmission electron microscopy. Protein levels of LC3, ATG5, CyclinD1, Bcl-2, Caspase-3, and c-Myc were determined by Western blotting. CCK8 assay and Transwell assays were used to determine cell viability and migration rate of HEp-2 cells, respectively. Flow cytometry was performed to analyze the rate of apoptosis. Immunofluorescence was performed to determine the expression distribution of LC3.

**Results:** Glut-1 knockdown significantly promoted autophagosome formation by upregulating the ratio of LC3-II/LC3-I as well as the role of rapamycin (RAP) and Beclin-1 overexpression on autophagy flux in HEp-2 cells. Glut-1 inhibition also reduced the viability of HEp-2 cells followed by the decreases in expression of cyclinD1 and c-Myc. In addition, Glut-1 depletion increased the number of apoptotic HEp-2 cells accompanied by activation of caspase-3 and downregulation of Bcl-2. Glut-1 knockdown also reduced the migration rate of HEp-2 cells by promoting the expression of N-cadherin and inhibiting the expression of E-cadherin. Beclin-1 consumption significantly reversed Gult-1 knockdown-mediated autophagy activation, resulting in promotion of both proliferation and migration and inhibition of apoptosis.

**Conclusions:** Glut-1 knockdown-induced autophagy inhibits the proliferation and migration of HEp-2 cells, and promotes apoptosis of HEp-2 cells partly by regulating autophagy.

## Introduction

Laryngeal squamous cell carcinoma (LSCC), which is related to many risk factors, including smoking, drinking alcohol, and laryngeal reflux, is one of the most common malignant tumors occurring in the head and neck [1]. Some studies have demonstrated that autophagy and glucose metabolism play important roles in the occurrence and development of cancer cells[2–5]. However, the mechanisms underlying autophagy and glucose metabolism in the process of LSCC remain unclear.

It has gradually become apparent that most tumor cells show a variable glycolysis phenotype, so increased glycolytic metabolism has been identified as a characteristic of malignant cells, known as the Warburg effect [4]. Previous studies confirmed the upregulation of members of the glucose transporter family (Gluts) during the process of glycolysis in tumor cells to meet the energy needs of cancer cells [6–8]. Glut-1, as an important carrier of glucose uptake and a major protein involved in the transport of glucose, plays a key role in maintaining the homeostasis of the body[9]. It has been reported that Glut-1 transporters are expressed at higher levels in human solid cancer cells compared to their normal counterparts[10–12]. Glut-1 also affects cell proliferation and chemoradiotherapy sensitivity of head and neck squamous cell carcinoma, including laryngeal carcinoma13-18. Our previous studies suggested that Glut-1 may affect the proliferation, glucose uptake, chemosensitivity, and radiosensitivity[8,13–17]. Although we found that the mechanism may be mediated through the PI3K/Akt pathway, the role of the activated PI3K/Akt pathway in this process was not significant. Therefore, further studies are required to investigate the mechanism by which Glut-1 results in proliferation, migration, and invasiveness.

Autophagy through Glut-1 translocation has been shown to affect glucose uptake by cancer cells[19–23]. Autophagy is the process by which damaged proteins or organelles are degraded and recycled by eukaryotic cells, and it plays a dual role in promoting cell survival and inducing cell death during cancer growth and migration[19–23]. The energy supply of cancer cells in an anoxic environment is associated with high levels of Glut-1 expression and autophagy activation[24,25] suggesting that Glut-1-mediated energy supply and cell growth are involved in the process of autophagy.

Beclin-1 is a homolog of the yeast ATG6/Vps30 gene, which was first identified as a molecule mediating autophagy in mammals[26]. It plays a positive regulatory role in the process of autophagy[27,28]. During the process of autophagy, Beclin-1 and Class III PI3K kinase form a complex that promotes the formation of autophagic vacuoles [29]. Previous studies have shown that stable transfection of Beclin-1 promotes autophagy and reduces tumorigenicity, suggesting that Beclin-1-mediated autophagic cell death may be associated with inhibition of tumor cell growth [30]. Beclin-1 inhibition is closely associated with a poor prognosis in some cancers[31–33]. Conversely, some studies showed that Beclin-1 upregulation in the same type(s) of cancer led to activation of autophagy and increased clonogenic survival, indicating an association between Beclin-1-mediated autophagy flux and tumorigenesis[34,35]. Although our previous study showed high levels of Glut-1and Beclin-1 expression in head and neck cancer and a correlation between Glut-1 and Beclin-1 expression, it remains unclear whether Glut-1-mediated autophagy flux in the process of oncogenesis is associated with Beclin-1-induced activation of autophagy in laryngeal carcinoma.

In this study, we explored the relationship between autophagy activity and Glut-1 in laryngeal carcinoma HEp-2 cells and provided a theoretical basis for understanding the role of Glut-1 in the growth and migration of LSCC.

## Materials and methods

### Materials

Anti-CyclinD1 antibody was obtained from Proteintech Group (Rosemont, IL). Antibodies against c-Myc, Bcl-2, Caspase-3, Cleaved-Caspase-3, β-actin LC3, ATG5, and Beclin-1 were purchased from Abcam (Cambridge, MA). Rapamycin was obtained from Sigma (St. Louis, MO). Annexin V-FITC/PI was obtained from BestBio (Beijing, China). A CCK-8 kit was purchased from Dojindo (Kumamoto, Japan). PBS was obtained from Zsbio (Shandong, China). RPMI 1640 medium, FBS, and trypsin (0.25%) were purchased from Gibco (Grand Island, NY). Acrylamide and methylene were obtained from Amresco (Solon, OH). Penicillin-streptomycin, phosphatase inhibitors, and RIPA and BCA kits were obtained from Beyotime (Shenzhen, China). ECL substrate was purchased from Thermo Scientific (Waltham, MA). Protein marker was obtained from TransGen Biotech (Beijing, China). Horseradish peroxidase (HRP)-labeled goat anti-rabbit antibodies was obtained obtained from Boster BioTec (Wuhan, China). Mini-Protean Tetra cell and Transfer slot were obtained from Bio-Rad (Hercules, CA). Polyvinylidene difluoride (PVDF) membranes were obtained from Millipore (Billerica, MA).

### Cell culture

LSCC HEp-2 cells were cultured in RPMI 1640 medium supplemented with 10% fetal bovine serum (FBS) in an environment of 5% CO_2_ and 37°C in a cell incubator (Thermo Scientific). Medium was exchanged every 2 days. Cells in logarithmic growth phase were used for follow-up study.

### Vector construction and cell transfection

Specific interference fragments targeting the Glut-1 and Beclin-1 genes were designed and synthesized with reference to the human Glut-1 and Beclin-1 cDNA sequences in GenBank. A lentivirus vector with green fluorescent protein reporter group was provided by Shanghai Sangon Biotech (Shanghai). The interference fragment was inserted into the HIV-1 lentivirus vector and transfected into 293T cells to obtain virus supernatant. After testing the titer of the virus, it was used to infect the target cells. HEp-2 cells were added to 6-well plates at a concentration of 1.5×10^5^ cells/well. RPMI 1640 medium was added to each well to total volume of 2 ml. When the cells had grown to 50%–70%, 200 μl of virus supernatant solution was added for transfection. Transfection efficiency was observed under a fluorescence microscope at 72 and 96 hours after transfection.

For construction of the Beclin-1 overexpression vector, Beclin-1 cDNA was amplified by RT-PCR and inserted into the corresponding site of the eukaryotic expression vector, pcDNA3.1(+). Transfection assay was then carried out using Lipofectamine 2000 in accordance with the manufacturer’s instructions (Thermo Fisher Scientific, Waltham, MA).

### Transmission electron microscopy

After 48 hours of treatment with RAP or transfection with Glut-1 shRNA and lenti-Beclin-1, the cells were washed in PBS, fixed in 2.5% glutaraldehyde, postfixed in 1% osmium tetroxide, and gradually dehydrated in ethanol and acetone. After embedding with epoxy resin, they were cut into sections and stained with uranyl acetate and lead citrate. Autophagy of the cells was observed by transmission electron microscopy (TEM; Thermo Scientific).

### CCK-8 assay

Cells were inoculated into 96-well cell culture plates at 5000/well and cultured for 24, 48, or 72 hours. Then, 10 μl of CCK-8 (Dojindo) was added to each well and culture was continued for 2 hours. The fluorescence was determined spectrophotometrically at a wavelength of 450 nm using a microplate reader (Thermo Scientific).

### Flow cytometry

Cells were seeded into 96-well plates at 1×10^5^/well and grown in attachment culture for 48 hours. The cells were then treated with 0.25% trypsin (Gibco) and washed three times with precooled PBS (Zsbio). The cells were collected by centrifugation at 2000 rpm for 5 minutes. Aliquots of 10 μl of Annexin V-FITC solution (BestBio) and 5 μl of propidium iodide solution (PI) were added to 400 μl of 1×Binding Buffer and mixed. The cells were resuspended at a concentration of ~1×10^6^/ml with the mixed solution. After incubation at room temperature for 15 minutes in the dark, the fluorescence distribution was examined by flow cytometry (BD Biosciences, Franklin Lakes, NJ).

### Transwell assay

Suspensions of 5×10^4^ treated cells from each group in 500 μl of serum-free medium were inoculated into the upper chamber of a Transwell system for migration assay. Cells were added to the lower chamber of the Transwell system in 750 μl of RPMI 1640 medium containing 10% FBS for 8 hours. After the chamber was removed, the upper layer was wiped to pass through the cells. The cells were dyed with crystal violet and the number of perforated cells was determined under a microscope.

### Western blotting

Total protein was extracted with RIPA solution (Beyotime Biotechnology, Shanghai, China) and protein concentration was determined using a BCA kit. Protein samples were separated by SDS-PAGE. After blocking for 1 hour with 5% skim milk, the membranes were incubated with primary antibodies at 4°C overnight. The next day, the samples were incubated with HRP-conjugated goat anti-rabbit IgG (diluted 1:500) for 1 hour at 37°C. After washing three times with TBST (Tris-buffered saline and Polysorbate 20), a chemiluminescence system was used to measure the levels of specific proteins using an enhanced chemiluminescence (ECL) coloring solution.

### Quantitative real-time polymerase chain reaction (RT-PCR)

Total RNA was isolated according to the manufacturer’s instructions. Briefly, 1 μg of RNA was reverse transcribed using a First Strand cDNA Synthesis Kit (K1622; Fermentas, Burlington, ON, Canada) and polymerase chain reaction (PCR) using a SYBR Green qPCR kit (Merck, Darmstadt, Germany) with incubation at 37°C for 60 minutes, 85°C for 5 minutes, and 4°C for 5 minutes, followed by storage at −20°C. RNA primers were designed and synthesized by Sangon Biotech. Primers for CyclinD1, c-Myc, Bcl-2, N-cadherin, E-cadherin, and ATG5 are shown in **Table 1**. The 2^ΔΔCt^ method was used to calculate the relative expression levels of these genes.

### Immunofluorescence

After incubation for 48 hours, the cells was washed three times with PBS and fixed for 15 minutes at room temperature with 4% paraformaldehyde. Then, cells were treated with 0.2% Triton X-100 for 15 minutes and blocked using 5% BSA (SH30574.03; Hyclone, Logan, UT) for 1 hour at room temperature. The cells were subsequently incubated with anti-LC3 antibody (1:500) at 4°C overnight. The next day, the cells were scrubbed and washed three times with PBS for 5 minutes each time. Finally, cells were stained with secondary antibodies for 1 hour and subsequently with DAPI for 5 minutes, and examined by confocal laser scanning microscopy (LSM 800; Zeiss, Oberkochen, Germany).

### Statistical analysis

Each experiment was repeated at least three times. Data are presented as the mean ± standard deviation (SD) and were analyzed using SPSS 25.0 (SPSS Inc., Chicago, IL). The least significant difference (LSD) or Dunnett’s method in one-way analysis of variance ANOVA (LSD for variance and Dunnett’s for variance inequality) was used to analyze and compare the data between groups. In all analyses, *P*<0.05 and *P*<0.01 were taken to indicate significance and high significance, respectively.

## Results

### Glut-1 inhibition promotes autophagy in HEp-2 cells

To explore the role of Glut-1 in the autophagy process of laryngeal carcinoma, we first established stable Glut-1 knockdown cell lines. As shown in Fig. 1A, Glut-1 shRNA lentivirus was constructed successfully and showed very high efficiency of infection in HEp-2 cells (Fig. 1A). Western blotting and qRT-PCR experiments showed that Glut-1 protein and mRNA levels were significantly decreased in HEp-2 cells infected with Glut-1 shRNA lentivirus compared with control and negative control (NC) groups, indicating that Glut-1 was effectively knocked down in HEp-2 cells (Fig. 1B–1C). TEM experiments showed that there were few autophagosomes in the control and NC groups. However, the number of autophagosomes was markedly increased in HEp-2 cells transfected with Glut-1 shRNA, suggesting that Glut-1 may be involved in the autophagosome formation (Fig. 2A). As an autophagy activator, rapamycin (RAP) promotes the formation of autophagosomes. In addition, Beclin-1 upregulation also promoted the autophagy process. The data presented here indicated that the acceleration effect of Glut-1 inhibition on autophagosome formation was similar to the effects of RAP exposure and Beclin-1 upregulation in HEp-2 cells. In addition, Glut-1 depletion plus RAP and Glut-1 depletion plus Beclin-1 overexpression showed additive effects on autophagosome formation compared to cells exposed separately to Glut-1 shRNA, RAP, and lenti-Beclin-1 (Fig. 2A). The overexpression efficiency of Beclin-1 was confirmed by qRT-PCR (Fig. 3A–3B). Furthermore, the expression of autophagy-associated protein LC3 was observed in the Glut-1 shRNA group. In the Glut-1 shRNA group, the ratio of LC3-II/LC3-I was significantly increased as well as in RAP and Lenti-Beclin-1 exposure groups. In addition, the groups with combination of Glut-1 shRNA plus RAP and Glut-1 shRNA plus lenti-Beclin-1 showed higher LC3-II/LC3-I ratios (Fig. 2B). These results suggested that Glut-1 knockdown directly promoted autophagy in HEp-2 cells.

**Figure 1.**
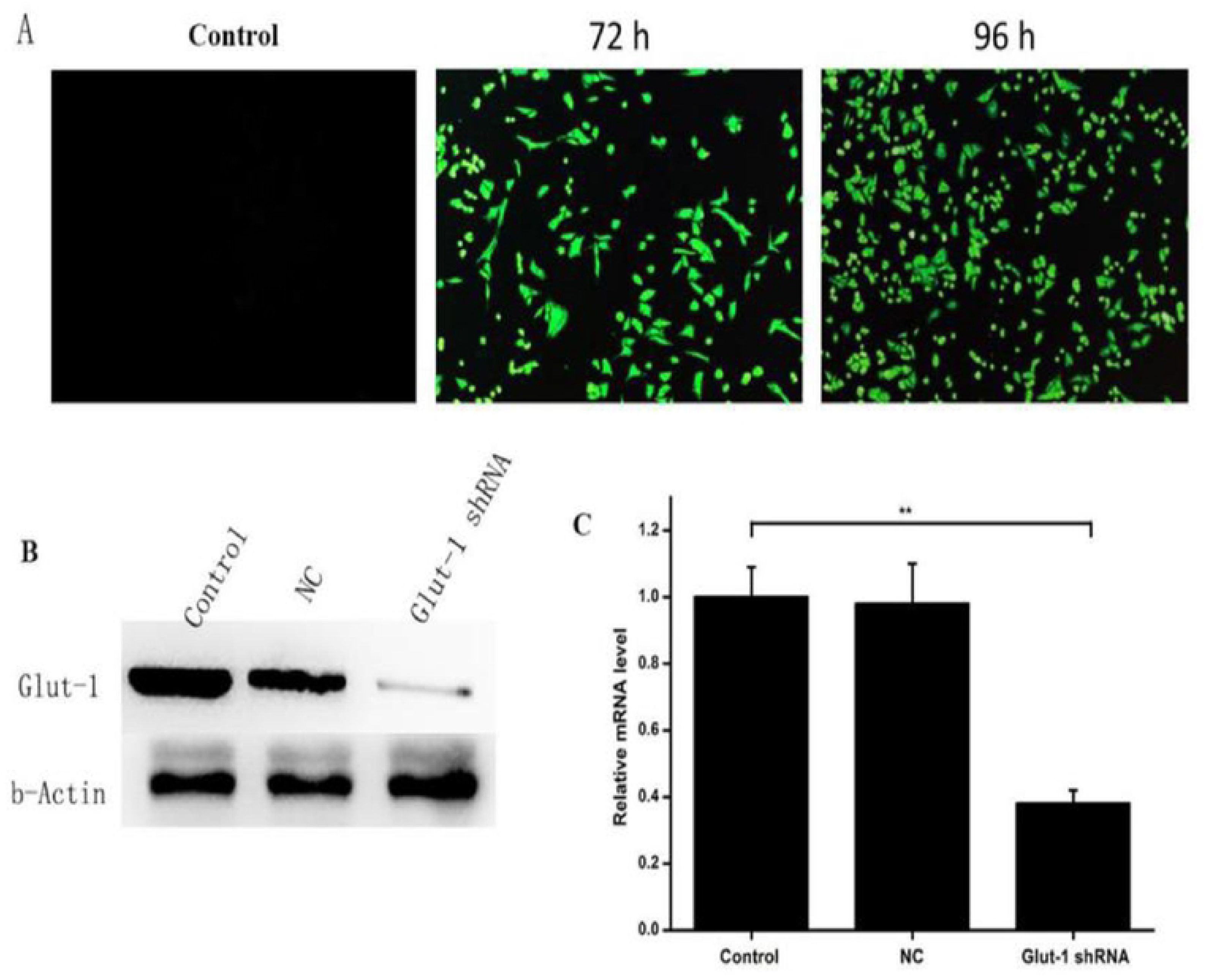
Construction of stable Glut-1 knockdown HEp-2 cells. **(A)** Infection efficiency of Glut-1 shRNA lentivirus observed by fluorescence microscopy. Green indicates infected cells. **(B)** The expression of Glut-1 protein in control, NC, and Glut-1 shRNA groups determined by Western blotting. **(C)** Relative mRNA expression of Glut-1 in the above groups determined by RT-PCR. Control, HEp-2 cells; NC, HEp-2 cells infected with empty lentivirus vector; Glut-1 shRNA, HEp-2 cells infected with Glut-1 shRNA lentivirus. Data are shown as the means ± SD, *n*≥3. \*\**P*<0.01.

**Figure 2.**
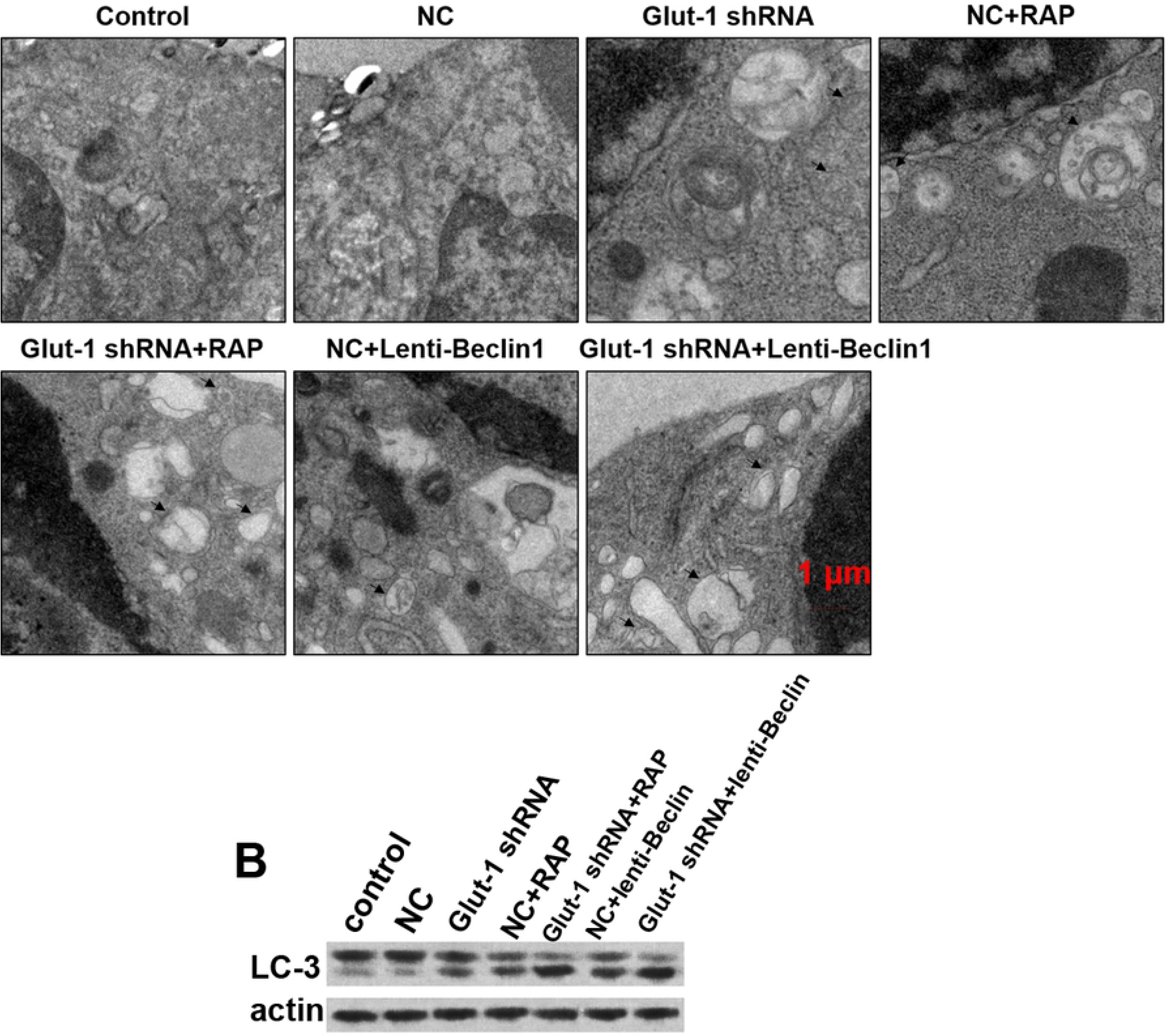
Effects of Glut-1 knockdown on autophagy in HEp-2 cells. **(A)** Transmission electron microscopy (TEM) images of HEp-2 cells in control, NC, Glut-1 shRNA, NC+RAP, Glut-1 shRNA+RAP, NC+lenti-Beclin-1, and lenti-Beclin-1 groups. N, nucleus; *, mitochondrion. The green arrow indicates autophagic body. The black arrow indicates autophagic lysosomes. **(B)** Expression of LC3 proteins in different groups determined by Western blotting. Data are shown as the means ± SD, *n*≥3. NC, negative control; Glut-1 shRNA, knockdown of Glut-1 in HEp-2 cells; RAP, rapamycin; lenti-Beclin-1, Beclin-1 overexpression plasmid.

**Figure 3.**
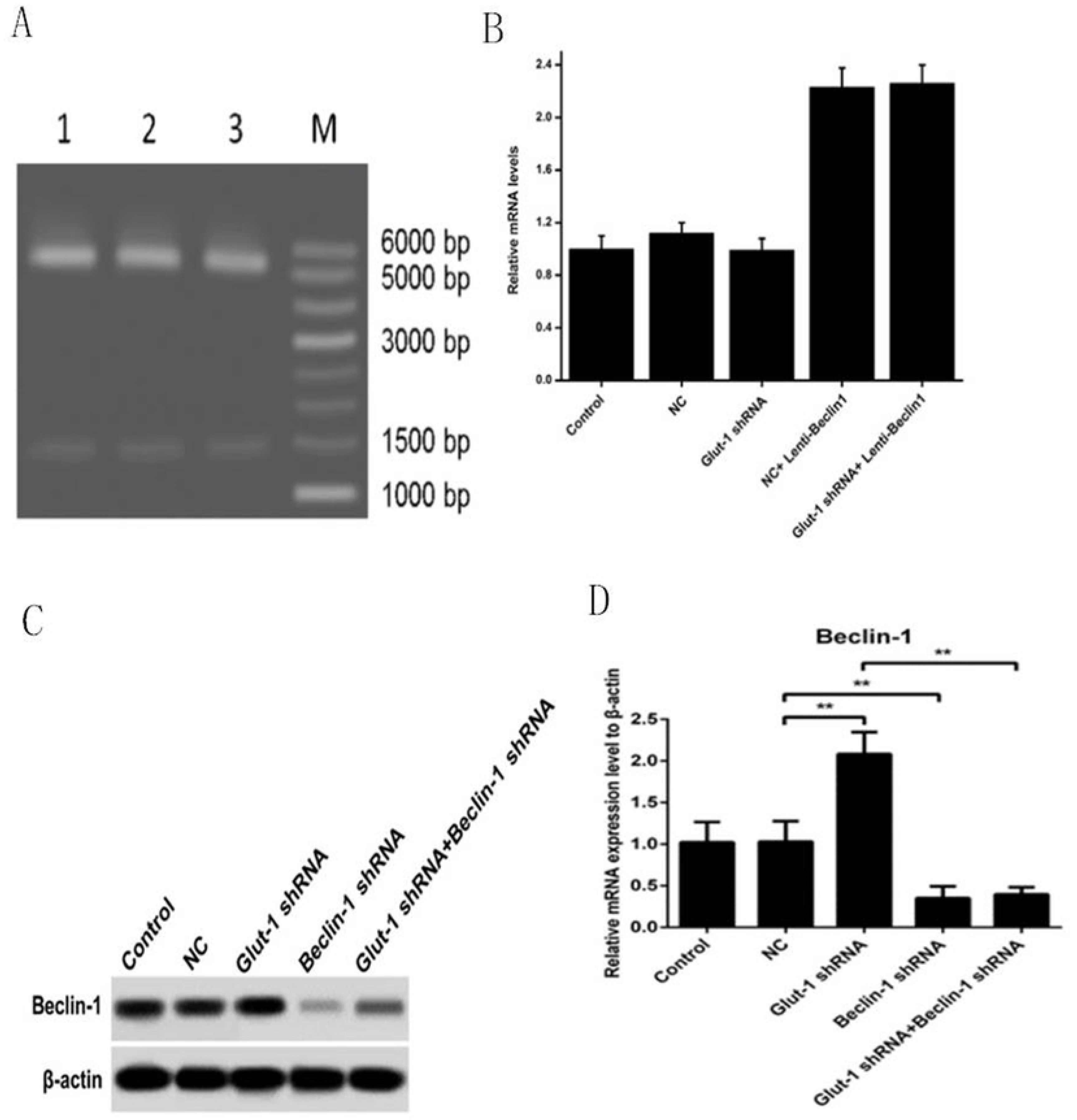
Construction of stable HEp-2 cell model infected with Beclin-1 shRNA lentivirus and overexpression plasmid. **(A)** Electrophoresis map of *Eco*RI and *Hin*dIII double digested plasmids. **(B)** Relative mRNA expression of Beclin-1 in control, NC, Glut-1 shRNA, NC+lenti-Beclin-1, and Glut-1 shRNA+lenti-Beclin-1 groups determined by RT-PCR. Lenti-Beclin-1, HEp-2 cells infected with Beclin-1 shRNA lentivirus. **(C)** Expression of Beclin-1 in control, NC, Glut-1 shRNA, Beclin-1shRNA, and Glut-1 shRNA+Beclin-1 shRNA groups determined by Western blotting. **(D)** Relative mRNA expression of Beclin-1 in different groups determined by RT-PCR. Beclin-1 shRNA, HEp-2 cells infected with Beclin-1 shRNA lentivirus. Data are shown as the means ± SD, *n*≥3. \*\**P*<0.01.

### Glut-1 consumption inhibits HEp-2 cell viability

Autophagy activation can inhibit the proliferation of tumor cells. As Glut-1 knockdown may be an inducer of autophagy, it may also affect the process of cell growth. The results of CCK8 assays indicated that both RAP treatment and Beclin-1 overexpression markedly inhibited the proliferation of HEp-2 cells. Importantly, knockdown of Glut-1 also significantly reduced the viability of HEp-2 cells at 24, 48, and 72 hours compared with control and NC groups, and addition of RAP and Lenti-Beclin-1 administration markedly increased the effect of Glut-1 depletion on proliferation inhibition (Fig. 4A). The levels of G1/S-specific CyclinD1 and c-Myc protein expression and relative mRNA expression were also decreased in cells with Glut-1 knockdown, RAP exposure, and lenti-Beclin-1 transfection separately. HEp-2 cells treated with Glut-1 shRNA plus RAP or Glut-1 shRNA plus lenti-Beclin-1 showed further reductions in the levels of cyclinD1 and c-Myc compared to individual treatments (Fig. 4B–4C). Therefore, our results suggested that Glut-1 knockdown-mediated autophagy may inhibit the proliferation of HEp-2 cells.

**Figure 4.**
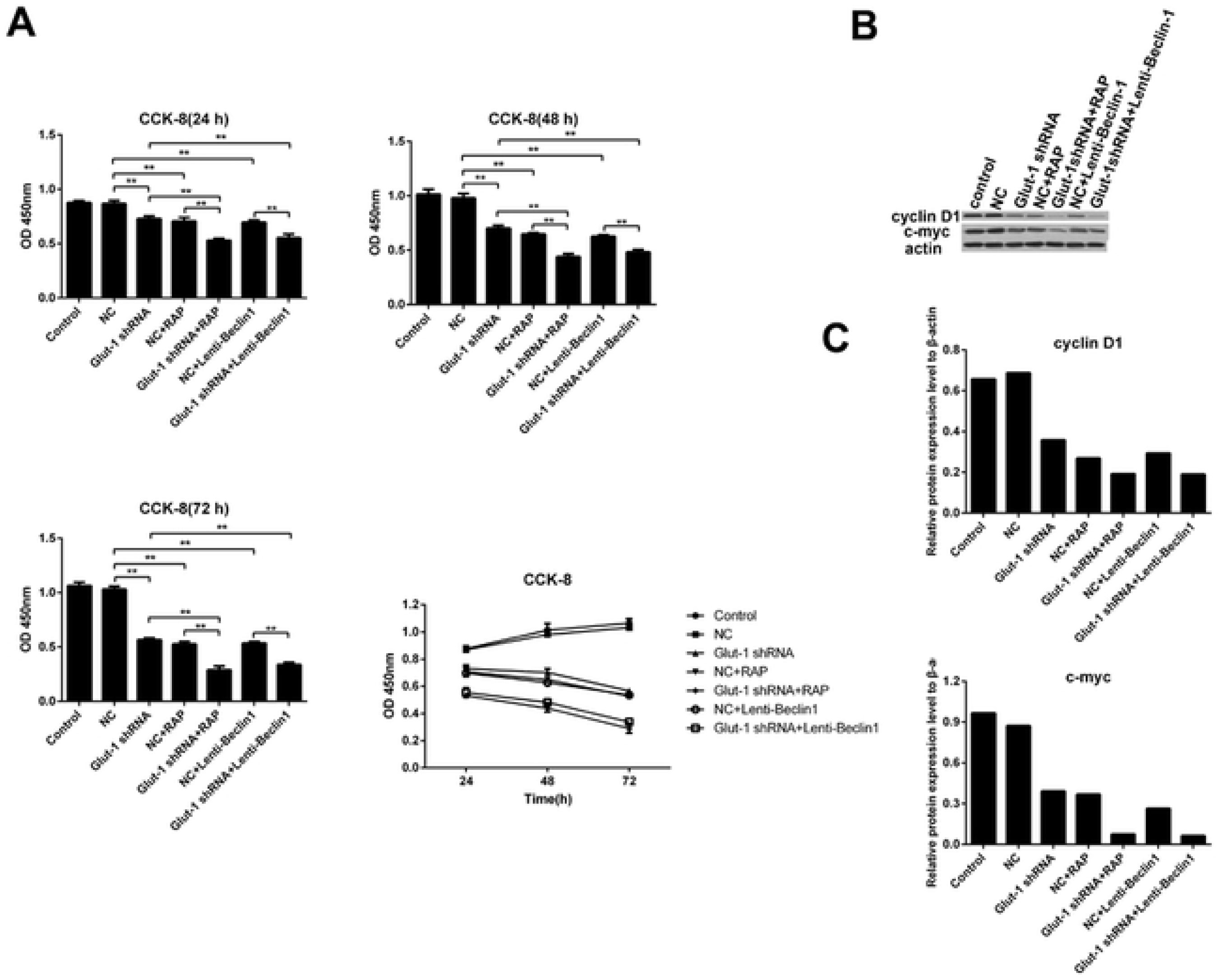
Impact of Glut-1 depletion on HEp-2 cell viability. **(A)** The proliferative activity of HEp-2 cells determined by CCK8 assay at 24, 48, and 72 hours in control, NC, Glut-1 shRNA, NC+RAP, Glut-1 shRNA+RAP, NC+lenti-Beclin-1, and lenti-Beclin-1 groups. **(B)** The levels of Cyclin D1 and c-Myc protein expression in the different groups determined by Western blotting. **(C)** Relative mRNA expression of Cyclin D1 and c-Myc in the different groups determined by RT-PCR. Data are shown as the means ± SD, *n*≥3. \**P*<0.05. \*\**P*<0.01.

### Glut-1 knockdown increases the apoptosis rate of HEp-2 cells

We further examined the effects of Glut-1 knockdown-mediated autophagy on cell apoptosis. The results of flow cytometry analyses suggested that Glut-1 knockdown significantly promoted the apoptosis rate of HEp-2 cells similar to the effects of RAP treatment and Beclin-1 upregulation. Greater numbers of apoptotic cells were observed in the Glut-1 inhibition plus RAP treatment group or Glut-1shRNA plus Beclin-1-overexpressing cells (Fig. 5A). As shown in Figure 3B, there were no significant differences in expression of caspase-3 between the groups. However, Glut-1 knockdown significantly upregulated the expression of cleaved-caspase-3 protein and downregulated the expression of anti-apoptotic Bcl-2 protein and the corresponding mRNA expression level of Bcl-2 compared with the control and NC groups (Fig. 5B–5C). The expression of cleaved-caspase-3 protein was further increased in the Glut-1 shRNA and RAP combination group and the Glut-1 shRNA and lenti-Beclin-1 combination group compared with the Glut-1 shRNA group (Fig. 5B–5C). The expression of Bcl-2 protein and corresponding mRNA expression in the Glut-1 shRNA plus RAP combination group and the Glut-1 shRNA plus lenti-Beclin-1 combination group were lower than in the Glut-1 shRNA group (Fig. 5B–5C). These observations confirmed that Glut-1 inhibition-induced autophagy enhanced the apoptosis of HEp-2 cells and showed an additive effect on cell apoptosis along with RAP or lenti-Beclin-1 treatment.

**Figure 5.**
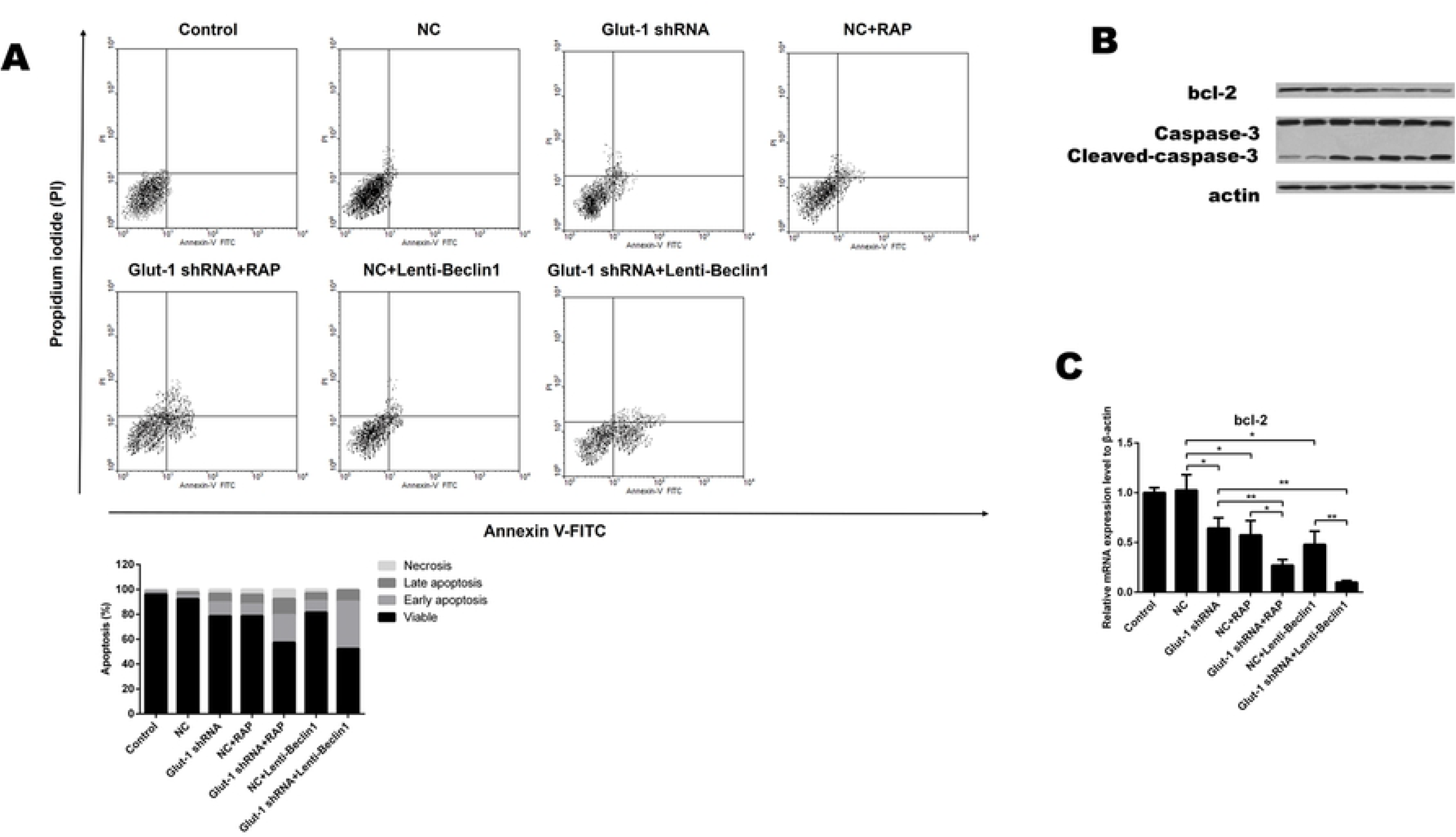
Role of Glut-1 downregulation on apoptotic rate of HEp-2 cells. **(A)** Total apoptosis rate in each group determined by flow cytometry using Annexin V/PI double staining. **(B)** Expression of Bcl-2, Caspase-3, and cleaved-caspase-3 proteins in different groups determined by Western blotting. **(C)** Relative mRNA level of Bcl-2 determined by RT-PCR. Data are shown as the means ± SD, *n*≥3. \**P*<0.05. \*\**P*<0.01.

### Inhibition of Glut-1 decreases migration ability of HEp-2 cells

Next, we evaluated the effects of Glut-1 inhibition on migration ability of HEp-2 cells. Transwell assay showed that Glut-1 knockdown significantly inhibited the number of migrating HEp-2 cells. The inhibitory effect of Glut-1 depletion on migration was markedly increased by combination with RAP treatment or Beclin-1 overexpression (Fig. 6A). Moreover, Glut-1 inhibition also downregulated N-cadherin protein and relative mRNA expression levels, but upregulated E-cadherin protein and relative mRNA expression compared with the control and NC groups (Fig. 6B–6C). The expression of N-cadherin was further decreased, while the expression of E-cadherin was significantly increased in the Glut-1 shRNA plus RAP group and the Glut-1 shRNA and lenti-Beclin-1 group compared with the Glut-1 knockdown alone group (Fig. 6B–6C). These results suggested that Glut-1 inhibition-evoked autophagy potentially increases the migration ability of HEp-2 cells.

**Figure 6.**
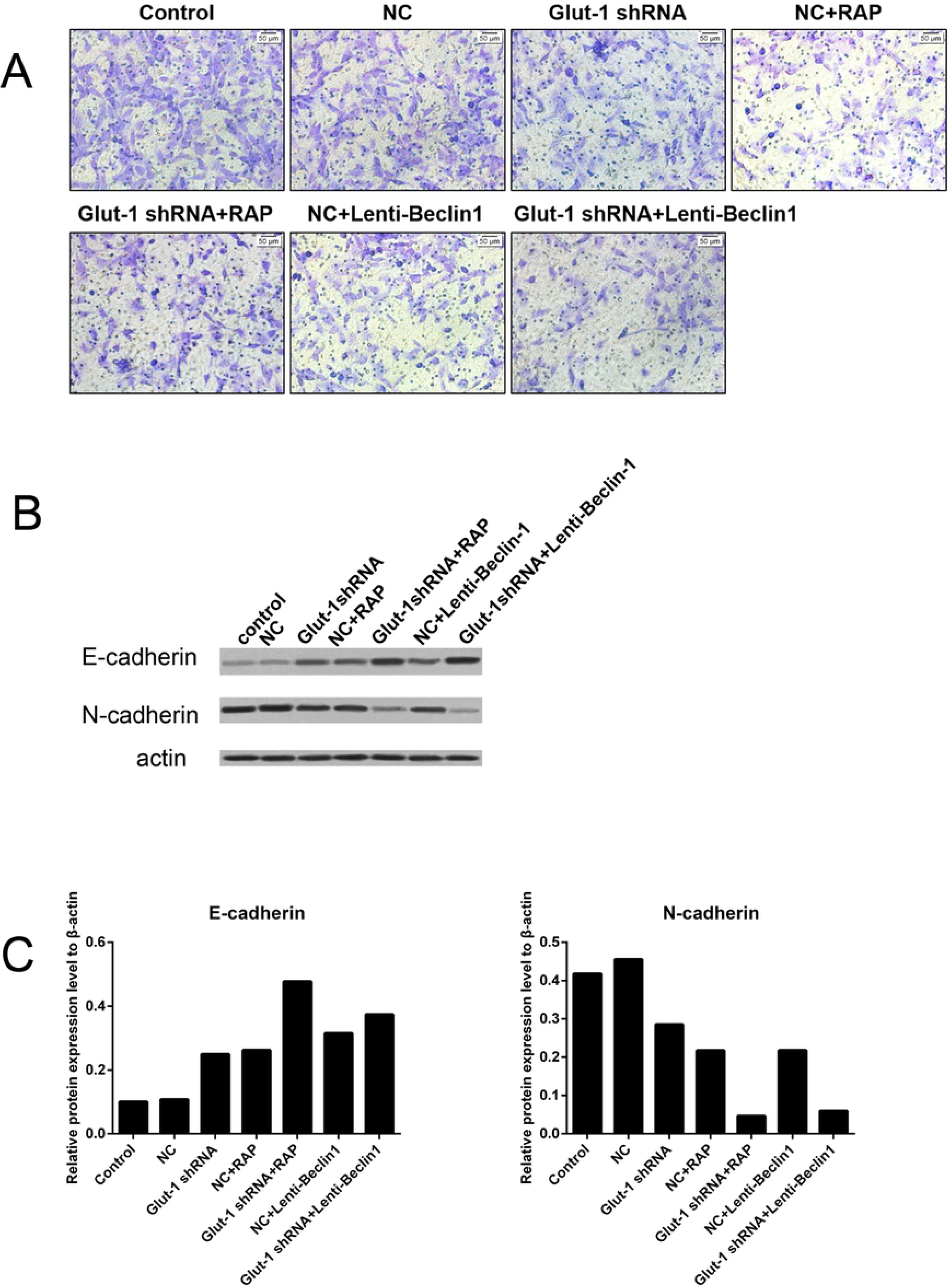
Effects of Glut-1 knockdown on migration ability of HEp-2 cells. **(A)** HEp-2 cell migration in each group determined by Transwell assay. **(B)** Expression of N-cadherin and E-cadherin in different groups determined by Western blotting. **(C)** Relative mRNA levels of N-cadherin and E-cadherin determined by RT-PCR. Data are shown as the means ± SD, *n*≥3. \**P*<0.05. \*\**P*<0.01.

### Knockdown of Beclin-1 rescues GLUT-1 inhibition-mediated autophagy

To investigate further the role of Beclin-1 in Glut-1 inhibition-induced autophagy activation, we established a stable HEp-2 cell line with Beclin-1 depletion (Fig. 3C–3D). As shown in Fig. 5A, Glut-1 knockdown markedly promoted, while Beclin-1 depletion significantly inhibited, the expression (protein and mRNA) of autophagy-related gene 5 (ATG5) and the ratio of LC3-II/LC3-I in HEp-2 cells (Fig. 7A-7B). Moreover, knockdown of Glut-1 significantly inhibited the expression of p62 protein, which was increased in cells transfected with Beclin-1 shRNA (Fig. 7A). However, the levels of ATG5 and LC3-II expression were significantly decreased, while that of p62 was restored in the Glut-1 shRNA and Beclin-1 shRNA combination group compared to the Glut-1 shRNA group (Fig. 7A–7B). Fluorescence staining also showed that Glut-1 knockdown promoted the expression of mRFP-LC3 and eGFP-LC3, while Beclin-1 inhibition reversed these phenomena and reduced their expression (Fig. 7C). These results suggest that Beclin-1 knockdown inhibited Glut-1 inhibition-mediated autophagy.

**Figure 7.**
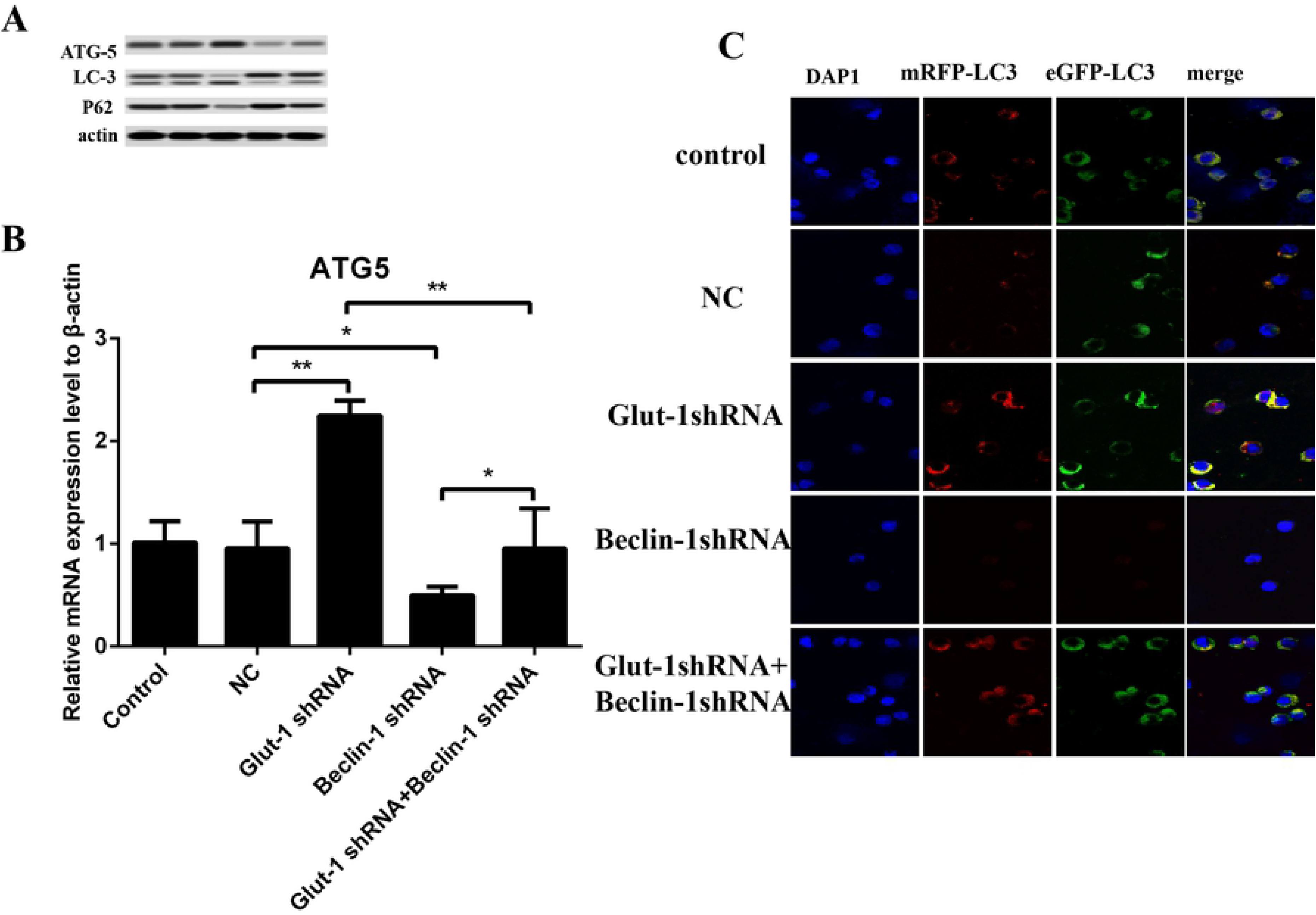
Effect of Beclin-1 on Glut-1 depletion-mediated autophagy. **(A)** Protein levels of ATG5, LC3, and p62 in control, NC, Glut-1 shRNA, Beclin-1 shRNA, and Glut-1 shRNA+Beclin-1 shRNA groups determined by Western blotting. **(B)** Relative mRNA expression of ATG5 in different groups determined by RT-PCR. **(C)** Expression of mRFP-LC3 and eGFP-LC3 in different groups determined by immunofluorescence analyses. Scale bar: 10 μm. Data are shown as the means ± SD, *n*≥3. \**P*<0.05. \*\**P*<0.01

### Beclin-1 is required in the process of Glut-1 knockdown-mediated improvement of tumor biological characteristics

We also measured the effects of Beclin-1 on Glut-1 knockdown-mediated proliferation inhibition. Our results showed that Beclin-1 knockdown significantly increased the viability of HEp-2 cells at 24, 48, and 72 hours. The Glut-1 depletion-mediated reduction of cell proliferation was significantly enhanced in cells exposed to Beclin-1 shRNA (Fig. 8A–8D). The results of flow cytometric analyses indicated that Glut-1 knockdown-mediated promotion on apoptosis of HEp-2 cells was blunted in the presence of Beclin-1 shRNA (Fig. 9A–9B). In addition, the inhibitory effect of Glut-1 knockdown on migration ability was also blocked by administration of Beclin-1 shRNA (Fig. 9C). However, both inhibition of cell apoptosis and migration and the promotion of cell proliferation capability were weaker in the Glut-1 shRNA plus Beclin-1 shRNA group than the Beclin-1 shRNA alone group. Taken together, these results confirmed that the Glut-1 knockdown-induced improvement of tumor biological characteristics is partly mediated by Beclin-1-induced autophagy activation.

**Figure 8.**
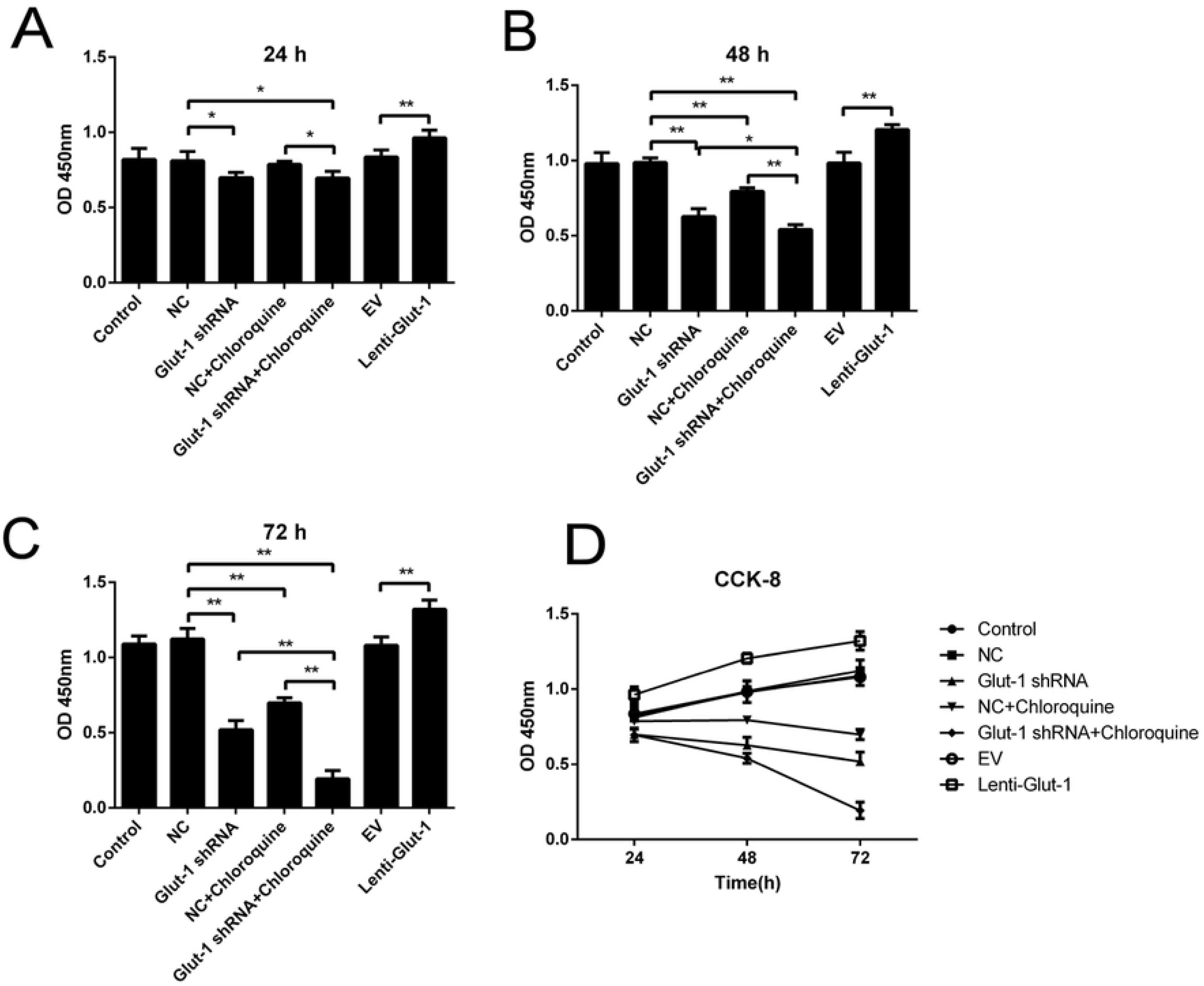
Beclin-1-mediated regulation of Glut-1 depletion-induced proliferation inhibition. The proliferative activity of HEp-2 cells was examined by CCK8 assay at 24 **(A)**, 48 **(B)**, and 72 hours **(C)** in different groups. **(D)** The results of CCK8 assays for all groups at all three time points are shown as a line chart. Data are shown as the means ± SD, *n*≥3. \**P*<0.05. \*\**P*<0.01.

**Figure 9.**
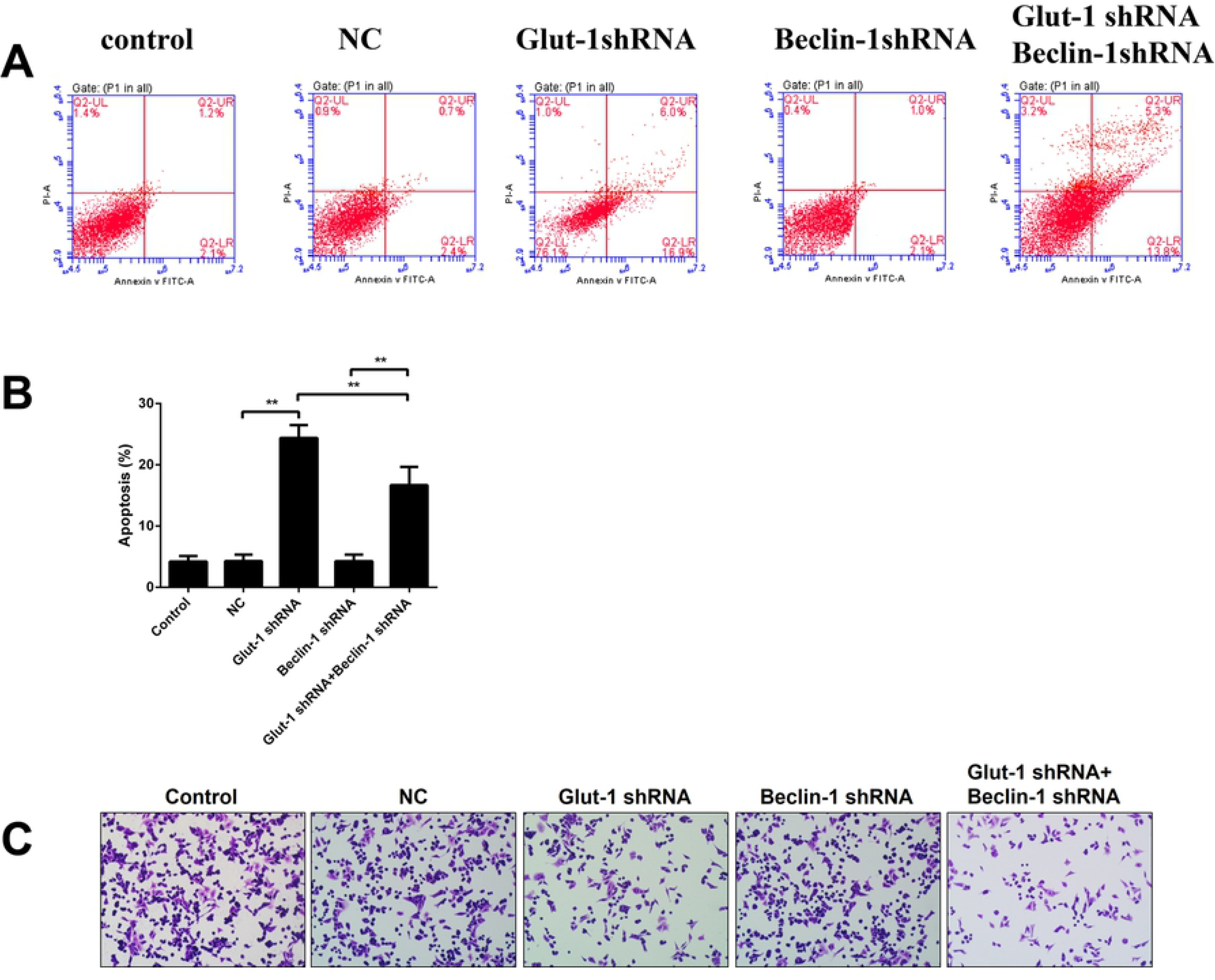
Effects of Beclin-1 on GLUT-1 depletion-mediated apoptosis promotion and migration inhibition. **(A)** Apoptosis in each group was determined by flow cytometry. **(B)** Relative apoptosis rate in each group. **(C)** Migration activity in each group determined by Transwell assay. Data are shown as the means ± SD, *n*≥3. \**P*<0.05. \*\**P*<0.01.

## Discussion

Autophagy is the process of capture, degradation, and recycling of organelles and proteins in lysosomes [36]. Previous studies indicated that autophagy activation has a dual effect on the occurrence of tumors [37]. In most cases, it was shown that activated autophagy of cancer cells can induce autophagic cell death, which may represent a promising approach for cancer treatment[19–31]. In the present study, we showed that Glut-1 knockdown evoked autophagy activity in HEp-2 cells leading to subsequent cell apoptosis, suggesting that Glut-1 may be a novel inducer of cell autophagy and useful for the treatment of laryngeal carcinoma.

Preliminary studies have shown that autophagy through Glut-1 translocation affects glucose uptake of cancer cells[19–23]. In comparison with normal cells, cancer cells preferentially produce energy by glycolysis in anaerobic environments [4]. The energy supply of cancer cells in hypoxic environments is related to the high-level expression of Glut-1 and autophagy. Many studies have shown that inhibition of Glut-1 in malignant tumors can effectively limit malignant tumor growth by inhibiting cell proliferation and migration, inducing cell cycle arrest and apoptosis[38,39]. In our previous study, the detection of 2-fluoro-2-deoxyglucose (FDG) concentration in head and neck cancer was associated with increased Glut-1 expression [31]. We also showed previously that inhibition of Glut-1 using antisense oligonucleotides reduced glucose uptake and promoted cell death in HEp-2 cells 8. Another study showed that Beclin-1 expression was negatively correlated with HIF-1α and Glut-1 expression and positively correlated with increased E-cadherin expression in human gastric cancer tissues, suggesting that autophagy deficiency promotes glycolysis and metastasis of gastric cancer in patients [40]. These studies suggested that Glut-1-mediated energy supply acts as a key regulator of tumor growth by affecting autophagy. Here, following knockdown of Glut-1 in HEp-2 cells, autophagy flux was significantly activated resulting in proliferation and migration inhibition and promotion of apoptosis compared to control cells. Based on the effect of autophagy on tumor cell death, we postulated that Glut-1 depletion-mediated autophagy activity was responsible for the death of HEp-2 cells, suggesting that targeting Glut-1 may be useful in the treatment of laryngeal carcinoma by regulating autophagy.

The mTOR (mechanical target of rapamycin) signaling pathway is considered to be one of the most important pathways regulating autophagy [41]. Rapamycin (RAP) can relieve the inhibition of ATG1 and promote autophagy by inhibiting the activity of mTOR42. Furthermore, activated ATG1 can form a protein complex through a series of ubiquitination-like ligation reactions with other autophagy-related factors, such as ATG3 and ATG5, in a manner dependent on the activity of phosphorylated Beclin-1[43]. Beclin-1 is a key protein in the process of autophagy in mammalian cells and is a direct mediator of autophagy[20,28,29,43]. Beclin-1 protein complex mediates autophagosome formation at the start of autophagy[20,28,29,43]. In this process, the free form of LC3 (LC3-I) in the cytoplasm forms LC3-II by binding to the phosphatidylethanolamine group and integrates into the autologous bilayer membrane structure to complete closure of the autophagosome[20,28,29,43]. In addition, the level of autophagy-related gene Beclin-1 expression in head and neck cancer tissues was shown to be significantly lower than that in adjacent tissues, indicating that Beclin-1-mediated autophagy participates in the progress of tumor formation [20]. In the present study, RAP markedly induced autophagy in HEp-2 cells. We also confirmed that Beclin-1 upregulation promoted, while Beclin-1 downregulation inhibited, autophagy followed by alteration of LC3-II/LC3-I rate, ATG5 level, and tumor characteristics, suggesting that Beclin-1 is critical for autophagy flux in laryngeal carcinoma development. Furthermore, Beclin-1 downregulation significantly restored the promotion effect of Glut-1 knockdown on autophagy activation and improvement of oncogenesis. However, the inhibitory effect on tumor growth was stronger with Beclin-1 inhibition alone than with Glut-1 shRNA plus Beclin-1 shRNA, suggesting that the mechanism of Glut-1 depletion-induced inhibition of tumor growth may be related to Beclin-1-mediated autophagy activation.

## Conclusion

This study was performed to explore the relationship between Glut-1 knockdown and Beclin-1-mediated autophagy activation in HEp-2 cells. Gult-1 knockdown-induced autophagy downregulated the proliferation of HEp-2 cells by decreasing the expression of Cyclin D1 and c-Myc, promoted apoptosis by decreasing the expression of Bcl-2 and increasing the activity of cleaved-caspase-3, and blunted the cell migration capability by downregulating N-cadherin expression and promoting E-cadherin expression. In contrast, Beclin-1 inhibition markedly reversed the tumorigenesis of HEp-2 cells induced by Gult-1 knockdown.

## Acknowledgments

This work was supported by National Natural Science Foundation of China (No. 81372903), and Science and Technology Department of Zhejiang Province, China (No.2016C33144).

## Author contributions

Wen-Dong Wang and Shui-Hong Zhou designed and wrote the manuscript. Jin-Long Zhu and Yang-Yang Bao collected materials and reviewed literatures. Jun Fan do experiments.

